# Integrated transcriptomic and metabolite analyses link DMSP cleavage to acetate overflow in the coral symbiont *Endozoicomonas ruthgatesiae*

**DOI:** 10.64898/2026.06.12.731890

**Authors:** Ying-Chieh Chen, Jui-Hung Yen, Ting-Chang Hsu, Wan-Ting Liao, Hsin-Feng Chang, Chih-Ying Lu, Li-Rong Lin, Po-Shun Chuang, Sen-Lin Tang

**Affiliations:** Biodiversity Research Center, Academia Sinica, Taipei city, Taiwan; Institute of Marine Environment and Ecology, National Taiwan Ocean University, Keelung city, Taiwan; Precious Instrument Center of New Ocean Researcher 2, National Taiwan Ocean University, Keelung city, Taiwan; Taiwan Ocean Genome Center, National Taiwan Ocean University, Keelung city, Taiwan

**Author notes:** Co-corresponding authors, Corresponding authors and email address: Sen-Lin Tang; Po-Shun Chuang.

**Keywords:** *Endozoicomonas*, DMSP, Acetate overflow, Transcriptomics, Thermal stress

## Abstract

*Endozoicomonas*, a dominant bacterial symbiont of corals, degrades dimethylsulfoniopropionate (DMSP) and produces dimethylsulfide (DMS), a key compound in the global sulfur cycle. By contrast, the carbon backbone of DMSP has received far less attention. Using *Endozoicomonas ruthgatesiae* strain 8E as a model, we exposed cultures to DMSP at 25°C and 31°C, simulating an ambient condition and thermal stress, respectively. Using gas chromatography-mass spectrometry (GC-MS) to monitor DMS and acetate production, we observed a significant increase in acetate efflux from 8E during DMSP metabolism in a growth phase-dependent manner. This increase in acetate was not observed when the native compound was replaced with ^13^C-labeled DMSP, suggesting that the additional acetate originated from DMSP. Transcriptomic analysis revealed that DMSP exposure upregulated *dddD* expression and triggered a reconfiguration of 8E’s central carbon metabolism, with upregulation of genes associated with the glyoxylate shunt-associated genes, and down-regulation of the TCA cycle and the Pta-AckA pathway. These findings suggest that DMSP-derived acetate overflows when its assimilation into biomass is restricted. Notably, elevated temperature diminished DMSP cleavage activity of 8E, indicating thermal sensitivity of this bacterial metabolic activity. Together, our results indicate that exposure to DMSP redirects carbon flow in *Endozoicomonas*, resulting in overflow of DMSP-derived acetate.

**Impact statement:** This study provides the first evidence of acetate efflux by *Endozoicomonas ruthgatesiae* strain 8E during dimethylsulfoniopropionate cleavage and suggests that this acetate overflow results from a shift in central carbon metabolism. These findings reveal an overlooked carbon output of bacterial DMSP metabolism, with potential implications for carbon exchange within coral holobionts.

**Data summary:** Data generated in this study are available in the NCBI Sequence Read Archive (SRA) under BioProject PRJNA1466041.

## Introduction

Coral holobionts are symbiotic assemblages comprising a coral host and various microorganisms, including photosynthetic dinoflagellates, bacteria, archaea, cyanobacteria, viruses, and microeukaryotes. Among these symbionts, photosynthetic dinoflagellates and bacteria are the best studied (Bonacolta et al., 2023; Peixoto et al., 2017). Photosynthetic dinoflagellates of the family Symbiodiniaceae, residing in coral gastrodermal cells, are the primary agents of photosynthesis in corals, providing fixed carbon to holobionts (Peixoto et al., 2017). On the other hand, bacteria are the major symbiotic prokaryotes in corals (Huggett & Apprill, 2019) and contribute to overall health and resilience of coral holobionts (de Breuyn et al., 2025; Maire et al., 2023). Metabolic pathways reconstructed from genomic analyses reveal that coral-associated bacteria potentially produce amino acids and vitamins essential for their hosts and for dinoflagellates (Hochart et al., 2023; Robbins et al., 2019). However, metabolite evidence for such contributions remains scarce.

Dimethylsulfoniopropionate (DMSP) and its degradation product, dimethylsulfide (DMS), reveal how bacteria interact metabolically with other symbionts in coral holobionts (Raina et al., 2010). DMSP is an important molecule in the marine sulfur cycle, produced by dinoflagellates (Curson et al., 2018), macroalgae (REED, 1983), bacteria (Curson et al., 2017; Kuek et al., 2022) and even by certain animals such as corals (Raina et al., 2013). In addition to serving as a source of carbon and sulfur (Stefels, 2000), microbial DMSP may function as an antioxidant (Sunda et al., 2002), an osmolyte (VAIRAVAMURTHY et al., 1985), a hydrostatic pressure protectant (Zheng et al., 2020), a chemoattractant (Seymour et al., 2010), or a signaling molecule (Johnson et al., 2016). In the ocean, three microbial pathways of DMSP metabolism have been identified (Shaw et al., 2022). Demethylation accounts for approximately 80% of DMSP transformation in marine environments (Curson et al., 2011) and generates methylmercaptopropionate (MMPA) (Schuller et al., 2012), which is then converted to methanethiol (MeSH) in subsequent CoA-mediated reactions (Moran et al., 2012). This process retains both sulfur and carbon in the marine food web (Curson et al., 2011). Cleavage constitutes the second pathway, accounting for 2–20% of total DMSP degradation (Kiene & Linn, 2000), generating DMS in combination with either acryloyl-CoA, acrylate, or 3-hydroxypropionate (3-HP). The latter can then be further converted to acetate. Although DMS is a well-known antioxidant in marine algae (Sunda et al., 2002) and a cloud-forming agent once released into seawater and the atmosphere (Kiene et al., 2000), the metabolic fate of acrylate and acetate remains to be determined.

The third pathway, oxidation, produces dimethylsulfoxonium propionate (DMSOP) and dimethyl sulfoxide (DMSO) (Thume et al., 2018) and accounts for the remaining catabolism of DMSP (Shaw et al., 2022). Notably, DMSP cleavage increases under hypersaline, oxidative conditions, and high UV irradiation (Kuek et al., 2022; Wang et al., 2022), implying a relationship between DMSP cleavage and bacterial stress response.

Acetate is a key intermediate in DMSP metabolism and a biologically useful carbon source. In *Escherichia coli*, excess acetate is excreted into the environment, a phenomenon known as “acetate overflow” (Bernal et al., 2016). This process highlights the mobility of acetate and its potential role as a “cross-feeding” molecule in symbiotic systems, a phenomenon describing excretion of nutrients by one organism and consumption by another (Rakoff-Nahoum et al., 2016; Rios-Covian et al., 2015). Therefore, we hypothesize that acetate generated from DMSP in corals flows to other organisms in the holobiont when in excess. However, to the best of our understanding, no studies have so far tested the linkage between DMSP cleavage and acetate overflow.

The bacterial genus, *Endozoicomonas* (*Gammaproteobacteria*; *Oceanospirillales*; *Endozoicomonadaceae*) is prevalent in corals and is known for DMSP degradation via the cleavage pathway (Chiou et al., 2023). In addition to corals (Neave et al., 2017; Pike et al., 2013), *Endozoicomonas* can be found in bivalves (Li et al., 2022), sponges (Nishijima et al., 2013), and ascidians (Schreiber et al., 2016). In corals, symbiosis with *Endozoicomonas* is closely linked to homeostasis of the holobiont, making *Endozoicomonas* putatively beneficial for corals (Peixoto et al., 2017). Single-cell transcriptomic data from a recent study showed a significant transcriptional response in alga-hosting coral cells following inoculation with *E. acroporae* Acr-14 (Lu et al., 2026). However, there is no evidence for metabolic interactions between *Endozoicomonas* and other coral symbionts. *Endozoicomonas ruthgatesiae* strain 8E (8E hereafter), isolated from *Acropora* sp. in Kenting, Taiwan, cleaves DMSP using the lyase, DddD, generating DMS and 3-HP. Chiou *et al*. (2023) showed that DMSP (0.1 mM) exposure induces metabolism of 3-HP to acetate in 8E, and they identified the pathway for subsequent incorporation of acetate into the tricarboxylic acid (TCA) cycle (Chiou et al., 2023), suggesting that DMSP may serve as an additional carbon source for 8E. However, it was not determined whether all acetate derived from DMSP cleavage was utilized by the bacterium. In this study, using stable isotopes and gas chromatography–mass spectrometry (GC-MS), we demonstrated that 8E also excretes DMSP-derived acetate into the environment. Transcriptomic analysis further offers a holistic view of the genetic mechanism underlying this DMSP-driven acetate overflow. These findings reveal the relationship between DMSP metabolism and acetate overflow in the coral-dominant *Endozoicomonas*.

## Methods

### Endozoicomonas ruthgatesiae strain 8E culture

For each experiment in this study, *Endozoicomonas ruthgatesiae* strain 8E was isolated from *Acropora* sp. in Kenting, Taiwan (Chiou et al., 2023), and activated from -80°C storage in Modified Marine Broth ver.4 (MMBv4) (Ding et al., 2016), a nutrition-rich medium, and subsequently enriched twice in MMBv4. Bacteria in this study were grown at 25°C with 200-rpm shaking unless indicated otherwise. The bacterial 16S rRNA gene was amplified from the bacterial culture using 341F (5’-CCTACGGGAGGCAGCAG-3’) and 1492R (5’-TACGGYTACCTTGTTACGACTT-3’) primers to confirm the identity of the bacterium. Briefly, 8E cells were resuspended in 10 µL of double-distilled water and incubated at 95°C for 5 min to lyse cells, after which the lysis solution (1 µL) was subjected to PCR. PCR conditions were as follows: initial denaturation at 94°C for 5 min; 30 cycles of denaturation at 94°C for 30 s, annealing at 55°C for 30 s, and extension at 72°C for 90 s, with a final extension at 72°C for 10 min. The PCR product was submitted to Genomics (New Taipei City, Taiwan) for Sanger sequencing and the resulting sequence was BLAST-searched against the NCBI database to assure bacterial species identity.

### DMSP metabolism assay

Following enrichment, the MMBv4 medium was removed by centrifugation at 4500 × *g* for 5 min and bacterial cells were resuspended in a minimal medium, supplemented with 0.5% (w/v) glucose (as a carbon source) and 0.2% (w/v) casamino acid (as a nitrogen source), as a starter culture for the subsequent metabolism assay. To assess DMSP metabolism, the 8E starter culture was inoculated into 30 mL of the same minimal medium at an initial 600-nm optical density (OD_600_) of 0.08 in nine 300-mL sterile, gas-tight vials, each supplemented with DMSP (Merck Sigma-Aldrich; n=3), [1,2,3-^13^C_3_] DMSP (synthesized and confirmed as described in (Chiou et al., 2023); n=3), or no substrate (negative control; n=3) **(Fig.1)**. One extra vial was supplemented with the same minimal medium and DMSP without the 8E culture to serve as an abiotic control for monitoring non-biological degradation of DMSP. As glucose is a carbon source preferred by 8E and because its metabolism also generates acetate, DMSP and [1,2,3-^13^C_3_] DMSP were supplemented at 3 mM, a relatively high concentration, to enhance the signal of DMSP-derived acetate. The DMSP metabolism assay was performed at both 25°C and 31°C to test the effect of temperature on 8E’s DMSP cleavage activity. At 0 h, 5 h, and 9 h of the assay, 6 mL of 8E culture aliquots were sampled from each vial using 10-mL syringe. A portion of extracted culture (1 mL) was subjected to OD_600_ measurement to estimate the growth of 8E. Another portion (2 mL) of extracted culture was centrifuged at 4500 × *g* for 5 min to pellet bacterial cells. Cell-free supernatants were used for estimating DMSP and acetate concentrations in the medium, while bacterial cell pellets were stored at -80°C for subsequent transcriptomic analyses. Other collected samples were kept in reserve on ice until all measurements were completed. OD_600_ measurements in this study were conducted using a ScanDrop spectrophotometer (Analytik Jena AG, Germany). For metabolism assays at both temperatures, three replicates of 0 h samples were collected from the starter culture (OD-adjusted), while for 5 h and 9 h samples, three replicates of each treatment were collected from individual vials.

**Figure 1.**
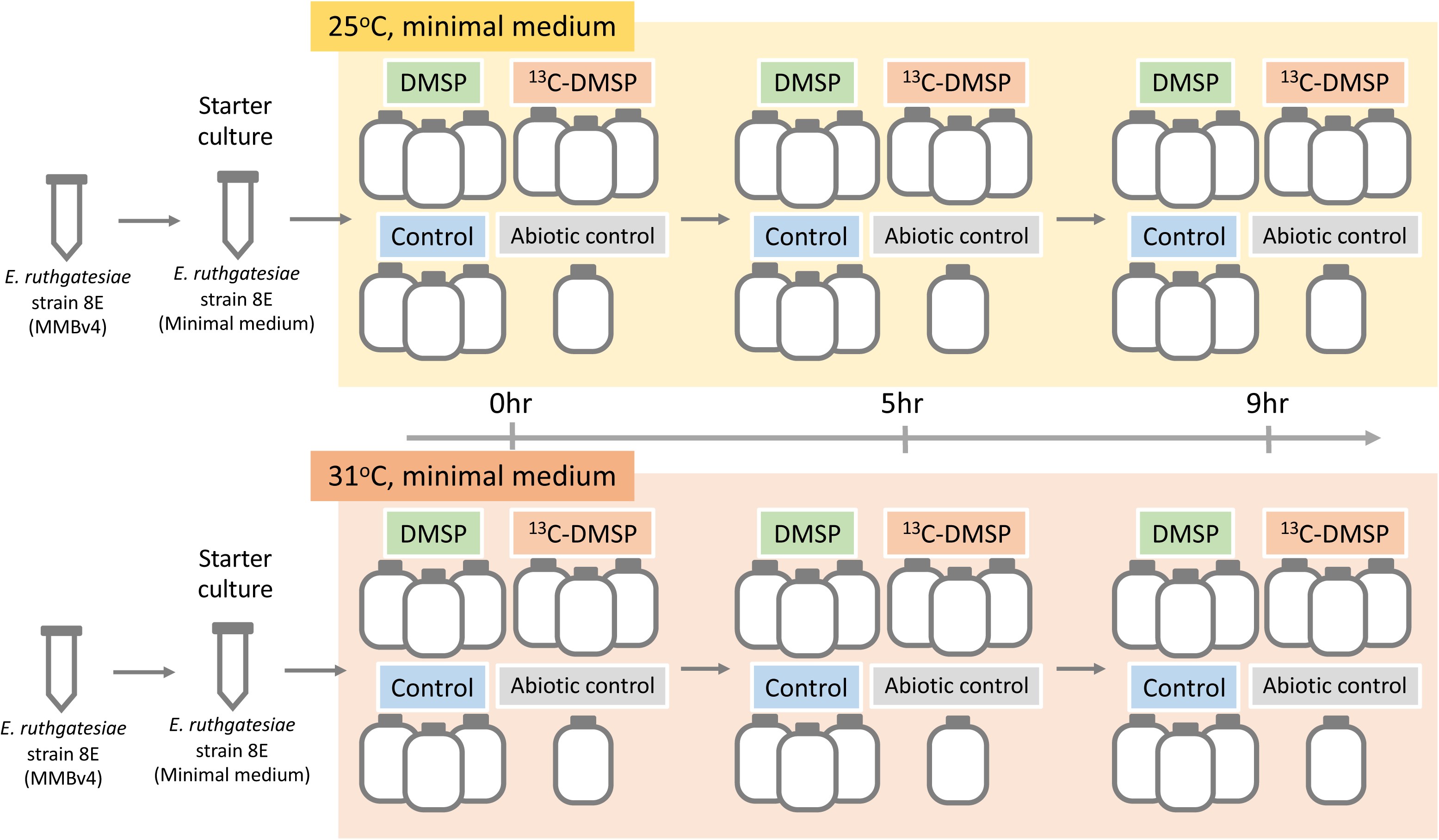
Group design in this study. The 8E culture in this research was enriched with MMBv4 (twice), changed into minimal medium, and three gas-tight vials were then inoculated before incubation prior to a metabolism assay and RNA sampling. All samples were incubated at 25°C and 31°C, respectively, and sampled after 0/5/9 h of incubation for DMSP/DMS/acetate/OD_600_ measurement.

### Growth curve and cell density measurement

To accommodate potential cell size differentiation among culture conditions, the growth curve and live cell density of 8E were measured using the same culture setup and conditions as used in the DMSP metabolism assay. The OD_600_ of the 8E culture at both 25°C and 31°C was measured at roughly 3-h intervals for 15 h. At 3 h, 6 h, and 8.5 h of incubation at both temperatures, 3 mL of the bacterial culture were collected. The minimal medium was removed by centrifugation at 3000 × *g* for 10 min and bacterial cells were resuspended in 3.5% NaCl and stained with LIVE/DEAD *Bac*Light Bacterial Viability Kits (Thermo Fisher Scientific, USA) for 15 min before counting using a CytoFLEX S Flow Cytometer (Beckman Coulter, USA). The linear relationship between OD_600_ and cell density was established for different culture conditions, and used for transforming OD_600_ measurements in the DMSP metabolism assay to live cell numbers. Growth curves and linear regressions between OD_600_ and 8E live cell density at different culture conditions are shown in **Fig. S1.**

### DMS Quantification

DMS production was quantified by collecting 500 µL of headspace air from the 300-mL vials using a 500-µL sampling syringe (Trajan Scientific, Australia). Gas samples were manually injected into an Agilent 6890N gas chromatograph system, equipped with an Agilent 19091S-433 model column (Agilent Technologies, USA). Samples were injected in split mode with a helium split ratio of 10:1. The inlet temperature was set at 200°C, with the injector pressure at 7.03 psi and the split flow at 14 mL/min. The GC-MS interface temperature was set at 325°C. Prior to each analysis, the column was thermally equilibrated for 0.5 min. Ion signals were detected using a 5975 Inert Mass Selective Detector (Agilent Technologies, USA). The signal at an m/z ratio of 62 ion was used for semi-quantification of DMS. The calibration curve for DMS quantification was established using the same vials at concentrations of 6, 3, 1, 0.5, 0.1, and 0.01 mM.

### Acetate Quantification

Acetate and U-^13^C-acetate (from ^13^C-DMSP) concentrations in the medium were determined by converting acetate into N-propyl-acetate following the method of Tumanov *et al*. (Tumanov et al., 2016). ^2^H_3_-acetate (Merck Sigma-Aldrich) was added to 1 mL of the supernatant at a final concentration of 0.2 mM as an internal standard. Derived samples were analyzed using the same GC-MS system as for DMS quantification, utilizing an Agilent 7683B Series automatic injector equipped with a 10-µL syringe (Agilent Technologies, USA). Samples were injected in split mode with a split ratio of 20:1. The inlet temperature was set at 280°C, the injector pressure at 7.3 psi, and the split flow at 24 mL/min. The GC-MS interface temperature was 325°C. Un-labeled acetate, U-^13^C-acetate, and the ^2^H_3_-acetate internal standard were quantified by integrating signal areas at corresponding m/z ratios (61 for un-labeled acetate, 63 for U-^13^C-acetate, and 64 for the internal standard). Signals of un-labeled and U-^13^C-acetate were subsequently normalized against the internal standard to calculate absolute acetate concentrations. Acetate concentrations were then normalized against live bacterial density to get the acetate production per bacterial cell. At each temperature and time point, differences in OD_600_, DMS, DMSP, and acetate concentration among treatments were assessed using one-way ANOVA followed by Tukey’s HSD post-hoc test.

### RNA extraction and transcriptomic data analysis

Bacterial cell pellets were resuspended in TRIzol reagent (Invitrogen, Thermo Fisher Scientific, USA) and RNA extraction was performed following the manufacturer’s instructions. RNA samples were submitted to BIOTOOLS (New Taipei City, Taiwan) for Next Generation Sequencing on a NovaSeq6000 system (paired-end; 150 bp x 150 bp). Resulting RNA-seq reads were quality-checked using FastQC (Andrews, 2010) and multiqc (Ewels et al., 2016) and processed with Trimmomatic (v0.39) to remove adapter sequences and low-quality bases (Bolger et al., 2014). Trimmed reads were then mapped to the published 8E genome (Chiou et al., 2023) using Hisat2 (Thakur, 2024), and gene expression was calculated using FeatureCounts (Liao et al., 2014). Gene counts were normalized using the median-of-ratios method and differential expression gene (DEG) analysis was conducted using DESeq2 (Love et al., 2014). Comparison was made between the DMSP and control groups at the same temperature and time point, as well as between samples of different temperatures, but the same treatment and timepoint. BlastKOALA (Kanehisa et al., 2016) was used for KEGG pathway mapping and pathway enrichment was analyzed using the *clusterProfiler* package in R. For statistical analysis, treatment and temperature were fixed at the whole plot level, whereas time was a repeatedly measured factor sampled from the same vial at multiple time points. As the standard PERMANOVA was not appropriate for testing all terms simultaneously, we performed PERMANOVA using a restricted, split-plot permutation design (Anderson & Braak, 2003), implemented with the *permute* package (Simpson, 2026) in conjunction with the *adonis2* function in the R package, vegan (Oksanen et al., 2026). Whole-plot terms (treatment, temperature, and their interaction) were tested by permuting whole vials, whereas split-plot terms (time and its interactions) were tested by permuting time points in each vial. Considering that transcriptomes of 0 h samples might reflect the shock of medium change, these samples were excluded from DEG analysis and PERMANOVA test.

## Results

### DMSP- and temperature-dependent growth of *E. ruthgatesiae* 8E

In this study we first tested growth of *E. ruthgatesiae* 8E exposed to 3 mM DMSP at 25°C and 31°C. While bacterial growth at 31°C was faster than that at 25°C, DMSP inhibited bacterial growth at both temperatures (**Fig. S1A,B**). The regression line of OD_600_ measurement and live cell density showed similar slopes in both DMSP-treated and control groups at both 25°C and 31°C. However, the intercept in the DMSP-treated group at 31°C was higher than that of the control group (**Fig. S1**). In an independent experiment, viable cells in DMSP-treated group consistently exhibited lower forward scatter (FSC) than those in control group at both temperatures between 3 and 8.5 h of incubation (**Fig. S1E,F**), suggesting that DMSP exposure reduced cell size of 8E. Cell densities in metabolism assays were thus calculated using regression equations in corresponding culture conditions.

In the metabolism assay at 25°C, OD_600_ values of the DMSP-(herein referred as the DMSP group) and [1,2,3-^13^C_3_] DMSP-treated groups (^13^C-DMSP group) were consistent in all conditions and timepoints (**Fig. 2A,B**). OD_600_ readings in the control group were significantly higher than in the DMSP and ^13^C-DMSP groups in both assays (*p*<0.05; **Table S1**). Live bacterial cell densities in the 25°C assay (**Fig. 2C**; calculated based on regression equations in **Fig. S1C**) paralleled the OD_600_ pattern at both 5 h and 9 h, with the control group showing higher density than the DMSP and ^13^C-DMSP groups. In contrast, calculated live cell densities in the 31°C assay exhibited no difference among treatments (**Fig. 2D**; based on regression equations in **Fig. S1D**), unlike the pattern of the OD_600_ measurement.

**Figure 2.**
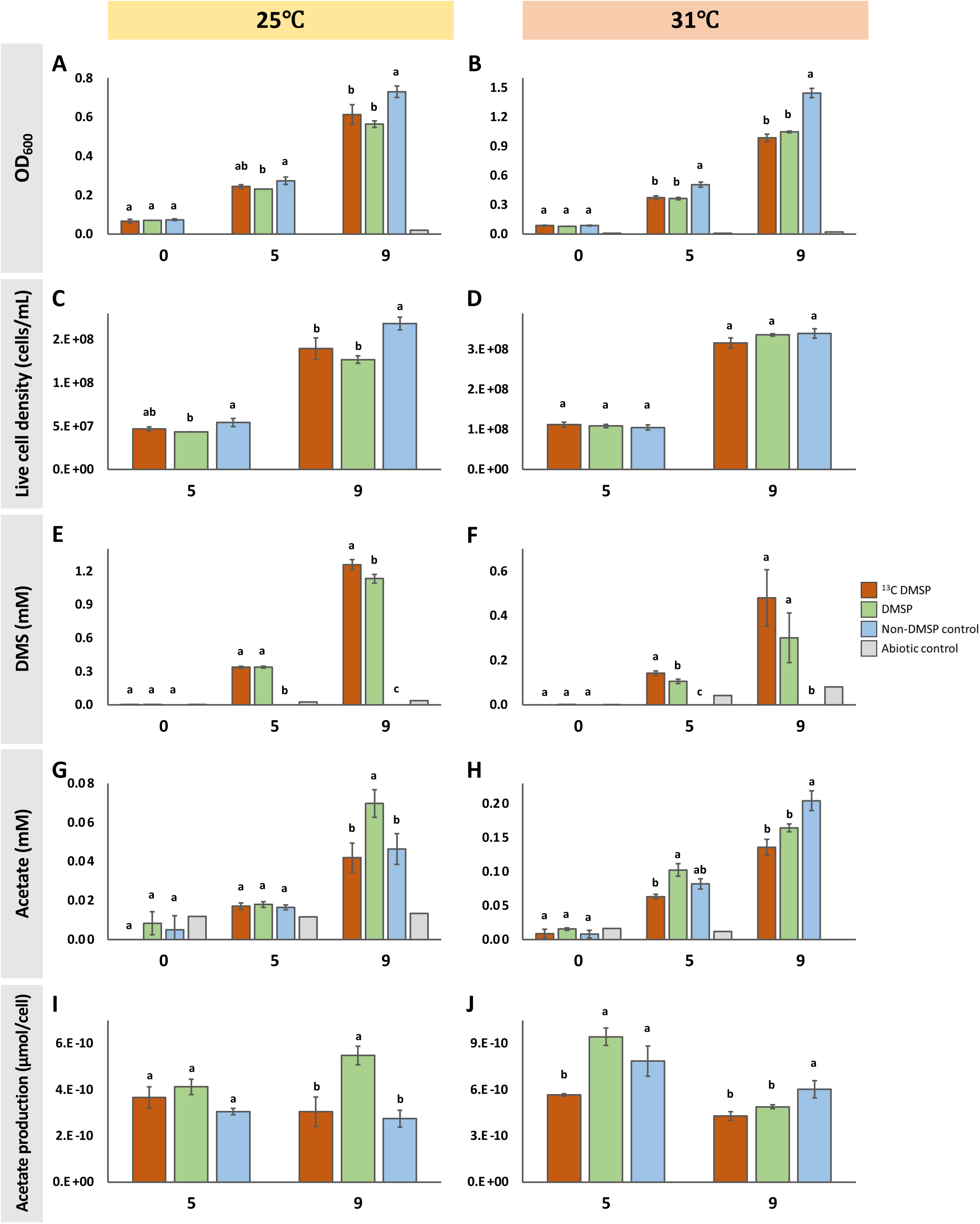
*Endozoicomonas ruthgatesiae* 8E cleaves DMSP to DMS and increases per-cell acetate release during DMSP exposure. (A,B) OD_600_ of *E. ruthgatesiae* strain 8E under 25°C and 31°C incubation. (C,D) OD_600_-converted live cell density, calculated using the regression line shown in Fig. S1C,D. (E,F) DMS production under 25°C and 31°C incubation. DMS was detected in the headspace of a 300-mL vial using GC-MS. (G,H) Acetate concentrations under 25°C and 31°C incubation. Acetate was detected in the culture medium using GC-MS. (I,J) Acetate production per cell at 25°C and 31°C. Data normalization was performed by dividing the detected acetate concentration by live cell density (cells mL ¹) of the sample. All measurements were performed in triplicate (n=3), except for the abiotic control group (n=1). Error bars represent standard deviation. No error bar is shown for the 8E negative control. Compact letter displays were used to denote statistically homogeneous groups having the same timepoint (α = 0.05, for ANOVA result see Table S1).

### *E. ruthgatesiae* 8E utilizes DMSP and releases acetate

DMS signals in the DMSP and ^13^C-DMSP groups increased in both metabolism assays (**Fig. 2E,F**). At 25°C, the ^13^C-DMSP group showed a DMS signal equivalent to that of the DMSP group at 5 h, but a significantly higher signal at 9 h. In the assay at 31°C, the DMS signal in the ^13^C-DMSP group was higher than that of the DMSP group at both 5 h and 9 h. However, the difference at 9 h was not statistically significant, due to great variance. At both 5 h and 9 h, DMS concentrations in the two DMSP-treated groups were higher at 25°C than at 31°C (**Fig. 2E,F**).

As with DMS production, acetate concentrations in the medium increased consistently in all treatments (**Fig. 2G,H**). However, acetate production per cell revealed different dynamics at 25°C and 31°C (**Fig. 2I, J**). In the 25°C assay, a significant increase in per-cell acetate production was detected in the DMSP group compared to the control at 9 h, whereas the level in the ^13^C-DMSP group remained comparable to that of the control. Meanwhile, the level of the DMSP group was higher at 9 h compared to at 5 h, whereas other groups showed no prominent difference between timepoints. In the 31°C assay, the per-cell acetate production in the DMSP group was higher than in the control and ^13^C-DMSP groups at 5 h, with the latter being statistically significant (*p*=0.0035). At 9 h, the control group was significantly higher than other groups. In contrast to the 25°C assay, all groups showed lower normalized acetate levels at 9 h than at 5 h. Despite the dynamic of un-labeled acetate in the medium, we detected no propyl-U-^13^C-acetate (m/z=63) in GC-MS in either the DMSP or ^13^C-DMSP group (data not shown).

### DMSP induces differential gene expression and metabolic pathway restructuring

In both metabolism assays, NovaSeq6000 RNA sequencing generated 0.8-1.3 Gbp of data per sample. Data processing yielded 6–10 million qualified, paired-end reads per sample, 84.0-88.2% of which were successfully aligned to the genome of 8E. No significant difference was found between 0-h samples in the two assays (PERMANOVA, R² = 0.525, *p* = 0.1; **Fig. S2**), indicating comparable baseline transcriptomes in both metabolic assays. Data from both assays were therefore analyzed together without further normalization. Principal component analysis (PCA) roughly separated data into 2 clusters: (1) the control group at 9 h in the 31°C assay and (2) all other samples (**Fig. 3**). At the whole-plot level, PERMANOVA showed that treatment (R² = 0.202, *p* = 4×10 ) and temperature (R² = 0.245, *p* = 1×10 ) significantly affected transcriptomic profiles, whereas their interaction was not significant (R² = 0.061, *p* = 0.138). At the split-plot level, incubation time (R² = 0.190, *p* = 0.0039) and its two-way interactions with treatment (R² = 0.056, *p* = 0.0039) and temperature (R² = 0.089, *p* = 0.0088) were all significant. The three-way interaction among treatment, temperature, and time was also significant (R² = 0.054, *p* = 0.033), indicating that the effect of DMSP treatment on 8E transcriptomes diverged between temperatures as incubation progressed **(Table 1)**. Consistent with this pattern, a sensitivity analysis testing the treatment-temperature interaction at each time point showed that the interaction was not significant at 5 h (*p* = 0.096), but became significant at 9 h (R² = 0.189, *p* = 5×10 ; **Table 2)**.

**Figure 3.**
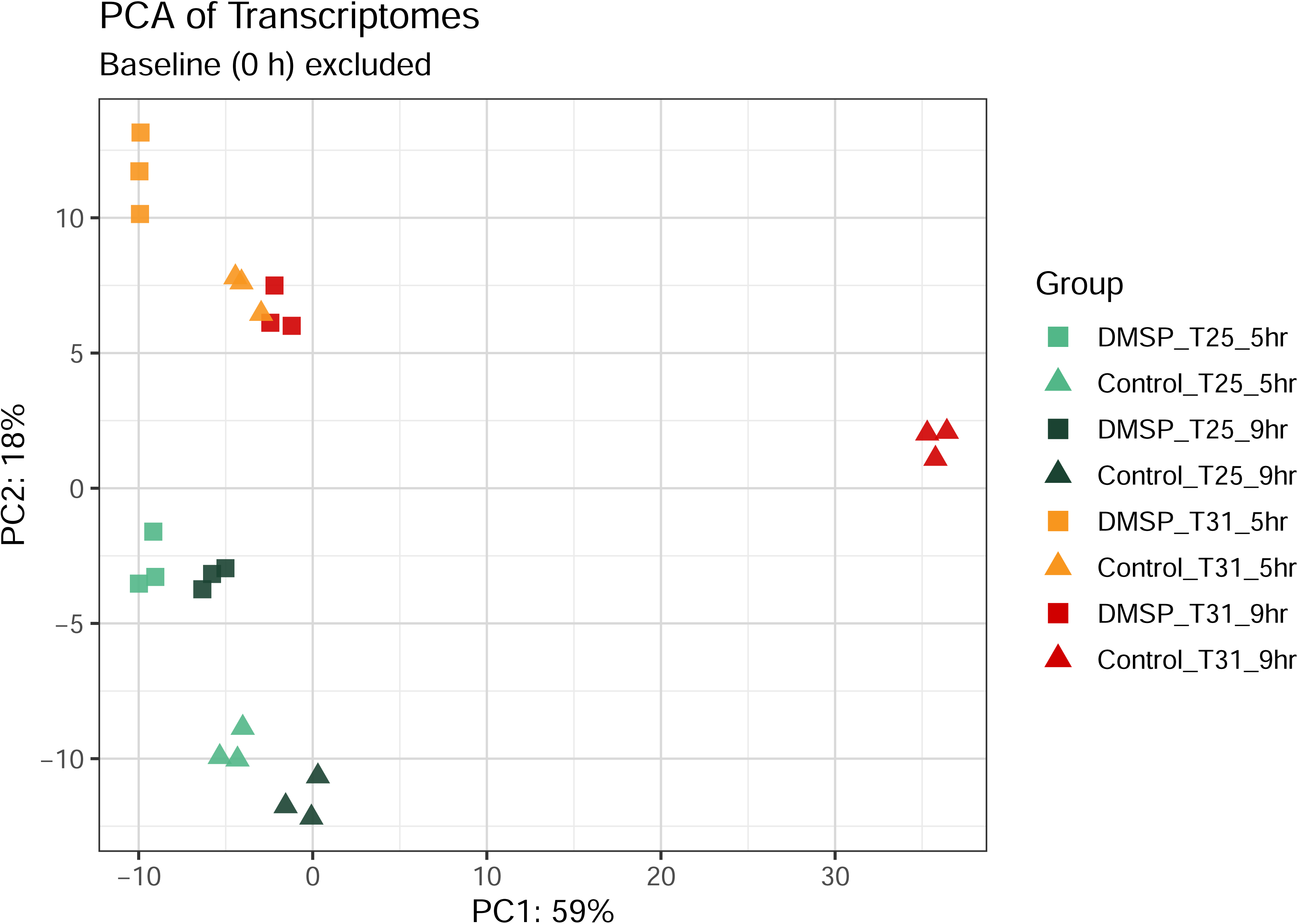
DMSP exposure shifts the global transcriptional profile of *Endozoicomonas ruthgatesiae* 8E at both 25°C and 31°C. Group differences in temperature, treatment, and time were assessed using split-plot restricted-permutation PERMANOVA on control- and DMSP-treated samples at 5 and 9 h (n = 3 independent vials [biological replicates] per treatment × temperature combination; see Table 1 and 2).

**Table 1.** Restricted-permutation (split-plot) PERMANOVA results for transcriptomic profiles.

| Error stratum | Term | R<sup>2</sup> | F | *p*-value |
|---|---|---|---|---|
| Whole-plot | Treatment | 0.2018 | 8.1903 | 0.0004 |
|  | Temperature | 0.2448 | 9.9365 | 0.0001 |
|  | Treatment × Temperature | 0.0606 | 2.4596 | 0.1375 |
| Split-plot | Time | 0.1897 | 28.8939 | 0.0039 |
|  | Treatment × Time | 0.0557 | 8.4853 | 0.0039 |
|  | Temperature × Time | 0.0886 | 13.4934 | 0.0088 |
|  | Treatment × Temperature × Time | 0.0536 | 8.1683 | 0.0332 |

Whole-plot terms were tested by permuting entire vials, treating each independently inoculated vial as the unit of replication (n = 3 per treatment × temperature combination). Split-plot terms were tested by permuting time-point labels in each vial. *p*-values were obtained via complete enumeration of the restricted permutation space.

**Table 2.** Sensitivity analysis: standard PERMANOVA performed independently at each time point.

| **Time** | **Term** | **R<sup>2</sup>** | **F** | ***p*-value** |
|---|---|---|---|---|
| 5 h | Treatment | 0.2514 | 11.5481 | 0.0003 |
|  | Temperature | 0.5189 | 23.8413 | 0.0001 |
|  | Treatment × Temperature | 0.0556 | 2.5537 | 0.0963 |
| 9 h | Treatment | 0.3552 | 27.1452 | 0.0001 |
|  | Temperature | 0.3511 | 26.8343 | 0.0001 |
|  | Treatment × Temperature | 0.1890 | 14.4450 | 0.0005 |

At each time point, every vial is represented exactly once. Accordingly, samples are independent and no permutation restriction is required.

At the gene level, differentially expressed genes (DEGs; *padj* < 0.05, |Log2 Fold Change| > 1) accounted for 1.6% and 1.1% of predicted genes in the 8E genome at 5 h and 9 h of the 25°C metabolism assay, respectively. More divergent proportions of DEGs were observed in the 31°C assay, ranging from 0.6% at 5 h to 8.2% at 9 h (**Fig. 4A**). However, only a small fraction of DEGs overlapped between different temperature-time combinations (**Fig. 4B,C**), with the majority of these overlapping genes annotated as hypothetical genes (**Table S2**). Among genes in the DMSP cleavage pathway, *dddD* exhibited strong up-regulation (*padj* < 0.05, |Log2 Fold Change| > 1) at all time points and temperatures (**Fig. 5B**). Moreover, *dddD* expression in DMSP-treated cultures was approximately two- to four-fold higher at 25°C than at 31°C. Compared to corresponding control groups, *dddD* expression in the DMSP-treated group was 14-79-fold higher in both metabolism assays. In addition to *dddD*, several genes in the DMSP cleavage pathway showed significant, but moderate up-regulation (*padj* < 0.05, |Log2 Fold Change| < 1), including: (1) *maeA* (up-regulation at both 5 and 9 h in the 25°C assay); (2) *glcB* and *icd2* (up-regulation at both 5 and 9 h in the 31°C assay); and (3) *maeB* (up-regulation in all conditions) (**Fig. 5**). On the other hand, *acs* and *bauC_2* showed moderate, but significant down-regulation in the 25°C assay (both at 5 and 9 h), and were strongly down-regulated at 9 h in the 31°C assay. Other genes in the pathway showed either slight down-regulation or no consistent regulatory pattern among conditions.

**Figure 4.**
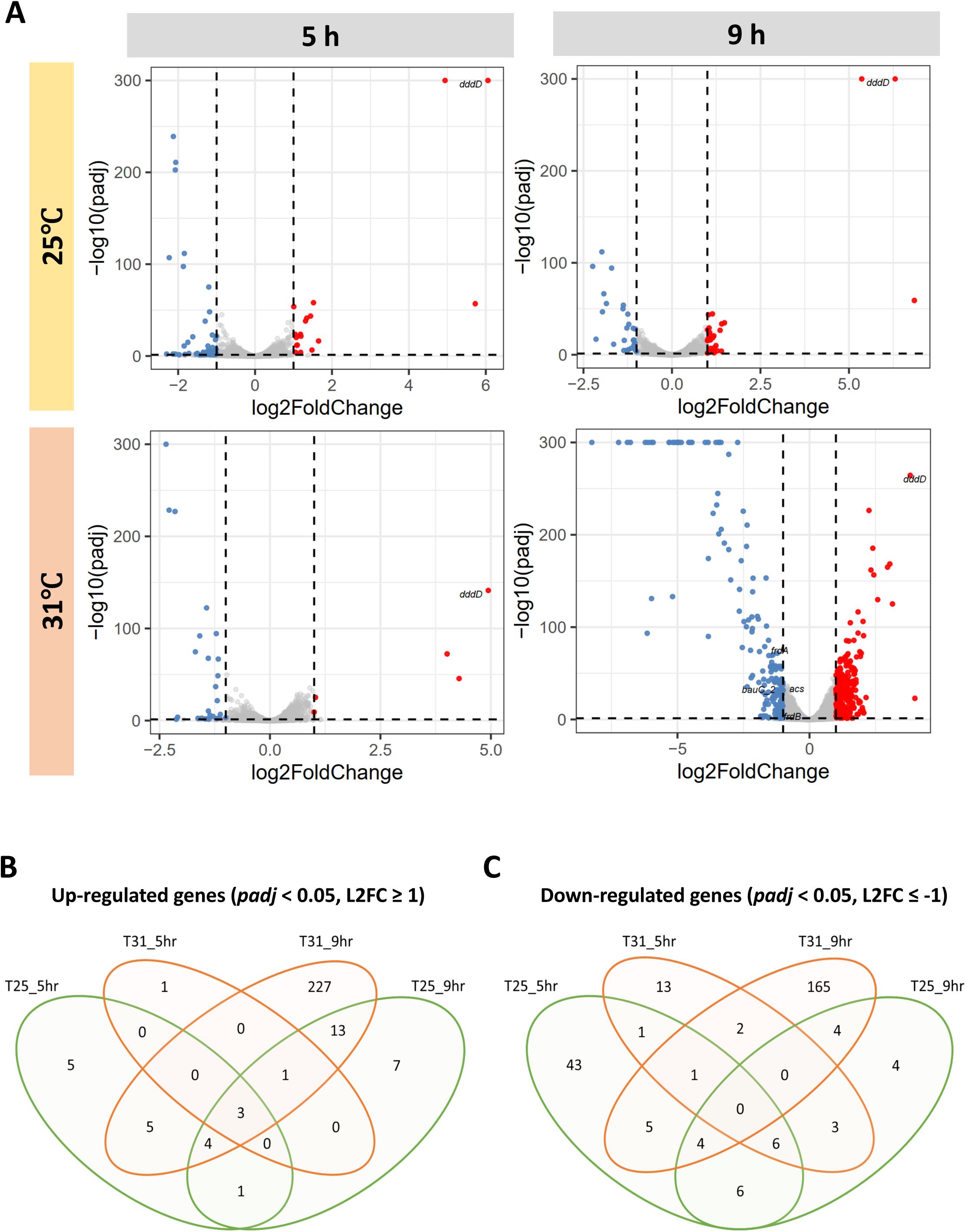
DMSP alters gene expression in *Endozoicomonas ruthgatesiae* 8E at both 25°C and 31°C. (A) Differentially expressed genes (DEGs) between DMSP-treated and control groups at different temperatures and time points. DEGs were identified as FDR < 0.05 and |log_2_(fold-change)| > 1. The *dddD* gene is labeled. (B,C) Venn diagrams of up- and down-regulated genes at different temperatures and time points. Each experimental condition was performed in triplicate (n=3).

**Figure 5.**
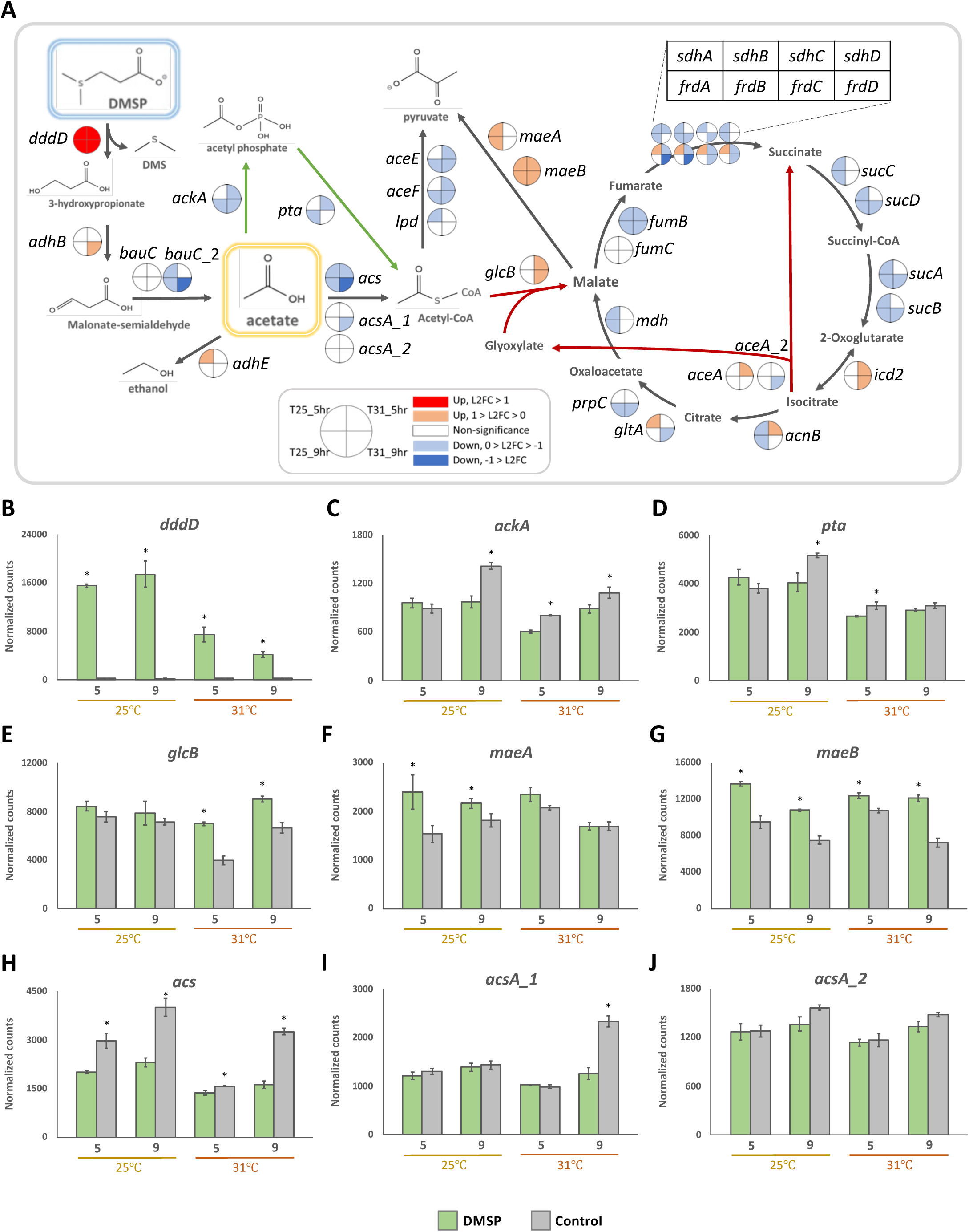
DMSP strongly induces *dddD* and alters expression of genes related to acetate- and central carbon metabolism in *Endozoicomonas ruthgatesiae* 8E. (A) Gene expression in the DMSP metabolic pathway. Circles along the pathway mark expression of corresponding genes, with colors indicating significantly different gene expression, estimated by DESeq2. Quarters of the circle representing different incubation times and temperatures. (B) *DddD* (C) *pta* (D) *ackA* (E) *glcB* (F) *maeA* (G) *maeB* (H) *acs* (I) *acsA_1* (J) *acsA_2* gene expression in DMSP-treated and control groups at 25°C and 31°C. Horizontal axis units from (B) to (J) are hours after incubation. 0 h samples are three technical replicates (n=3) collected from the same starter culture. For the 5 h and 9 h samples, three replicates of each treatment came from individual vials as biological replicates (n=3). Error bars represent standard deviation, and asterisks mark significant differences between the DMSP and control treatments in the same conditions (timepoint × temperature) (α = 0.05).

KEGG enrichment analysis revealed temperature-dependent responses (**Fig. 6**). At 25°C, pathways including the citrate cycle (TCA cycle), oxidative phosphorylation, the bacterial secretion system, glyoxylate and dicarboxylate metabolism, carbon metabolism, ABC transporters, lipoic acid metabolism, and other carbon fixation pathways were suppressed at 5 h, whereas biosynthesis of cofactors was the only significantly up-regulated pathway. At 9 h, suppression of the TCA cycle and oxidative phosphorylation continued, together with down-regulation of pathways for ribosome, 2-oxocarboxylic acid metabolism, valine, leucine, and isoleucine degradation, and microbial metabolism in diverse environments. No pathway showed significant activation at 9 h under 25°C. At 31°C, pathways significantly up-regulated at 5 h included biosynthesis of cofactors, cysteine and methionine metabolism, aminoacyl-tRNA biosynthesis, biotin metabolism, and homologous recombination, with no pathway being strongly suppressed (**Fig. 6**). At 9 h, quorum sensing, DNA replication, and homologous recombination were significantly up-regulated, whereas valine, leucine, and isoleucine degradation were strongly down-regulated.

**Figure 6.**
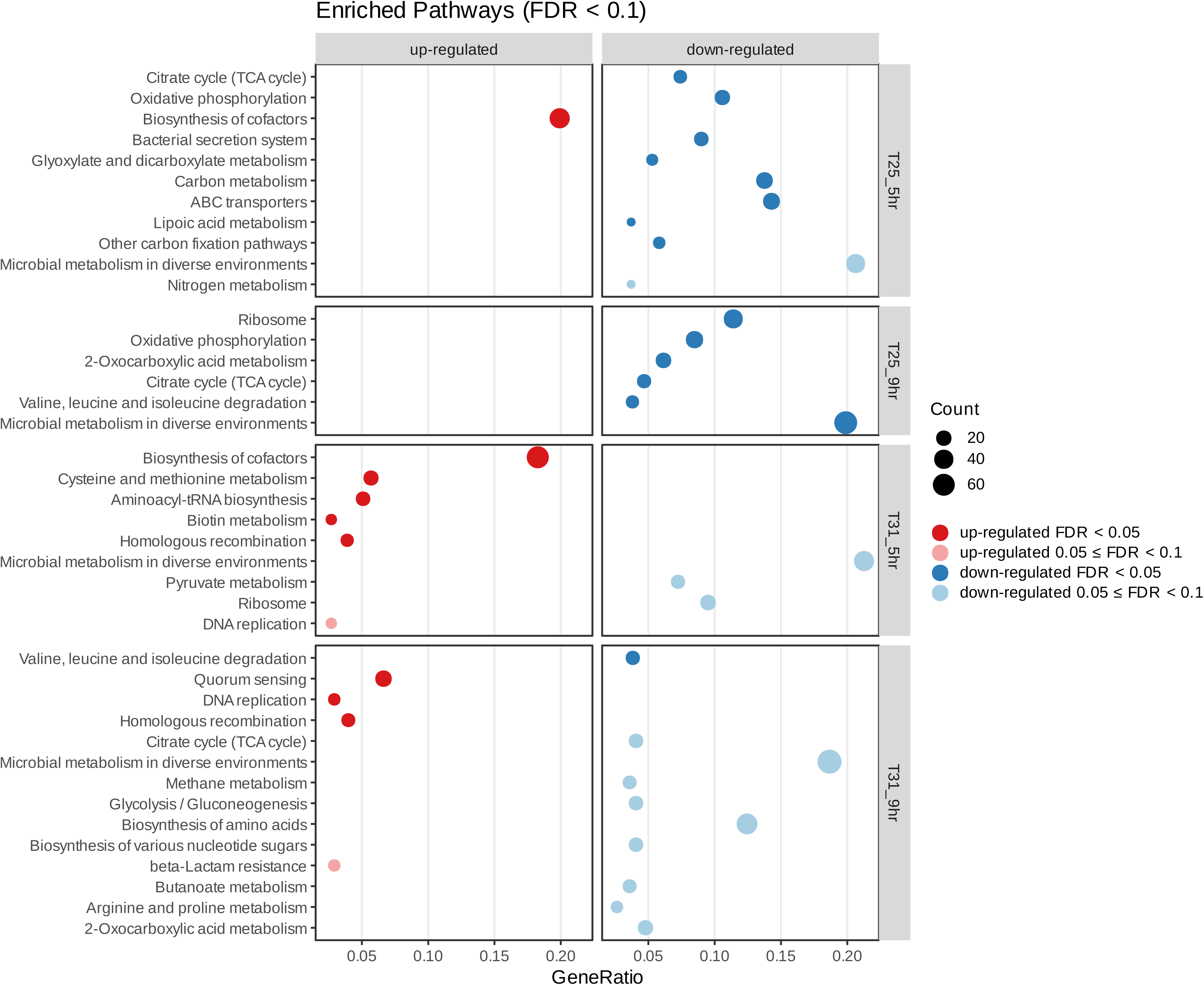
DMSP exposure alters metabolic pathways differently at 25°C and 31°C. Pathway enrichment was determined using the clusterProfiler package in R. Enriched pathways among up- and down-regulated genes under different FDR thresholds are marked with different colors. Pathways with FDR > 0.1 are not included.

## Discussion

*Endozoicomonas* bacteria catabolite DMSP into DMS and 3-hydroxypropionate (3-HP) though cleavage pathway. Chiou *et al*. (2023) identified upregulation of DMSP-degrading genes and those for converting 3-HP into acetate in *E. ruthgatesiae* strain 8E upon DMSP exposure (Chiou et al., 2023). Owing to its tendency for extracellular efflux and its metabolic utility (Bernal et al., 2016; El-Mansi & Holms, 1989; Pinhal et al., 2019), acetate represents a substrate for cross-feeding between organisms (Rios-Covian et al., 2015). In this study we employed 8E as a model and provide the first evidence of DMSP-driven acetate excretion in DMSP-degrading bacteria, both at an ambient coastal water temperature (25°C) and simulated thermal stress (31°C). The results of isotope-labeling experiments suggested that excreted acetate originates from carbon atoms in DMSP, and the transcriptomic analysis identified its putative genetic mechanism. These findings link the previously characterized DMSP cleavage capacity of *Endozoicomonas* to the extracellular release of DMSP-derived acetate, providing new insights into the potential ecological role of bacterial DMSP metabolism in coral holobionts.

### 8E excretes acetate during DMSP metabolism

DMSP metabolism activity was confirmed by *dddD* expression (**Fig. 5B**) and DMS production (**Fig. 2E,F**). We normalized acetate concentrations in the medium (mM) by live cell density (cells mL^-1^), representing acetate production per cell of 8E (µmol cell^-1^) during DMSP exposure (**Fig. 2I,J**). This reveals higher per-cell acetate excretion in the DMSP-treated group than in the control group at 25°C, 9 h. These results suggest a link between DMSP catabolism and acetate excretion in specific conditions. Bacterial culture at these conditions was at the early exponential (log) phase (**Fig, S1A,B**). Thus, the observed DMSP-driven acetate excretion may reflect growth stage-dependent acetate transport in 8E.

Both DMSP and glucose were provided in metabolism assays. This experimental design is essential, as 8E grows very slowly in glucose-free minimal medium, such that acetate excretion was undetectable (data not shown). This design, however, results in two sources of acetate in the medium. To address this issue, ^13^C-labeled DMSP was employed in our metabolism assays. A corresponding diminishment of increased unlabeled acetate was found in the ^13^C-DMSP-treated group at 25°C, 9 h and 31°C, 5 h (compared to the DMSP-treated group; **Fig. 2G,H**). This suggests that additional acetate excreted under DMSP exposure originated mostly from DMSP rather than from glucose in the medium. However, in this study we detected neither U-^13^C-acetate in the medium (GC-MS) nor ^13^C enrichment in 8E cells (Nano-SIMS; data not shown). These negative results may reflect the subtlety of the isotope signal and technical limitations inherent to detection methods. For example, sample preparation procedures for NanoSIMS such as fixation and washing have been reported to attenuate isotope enrichment by up to 80%, possibly leading to false-negative results (Meyer et al., 2021). Future studies employing optimized analytical strategies, such as Isotope Ratio Mass Spectrometry (IRMS) for higher-sensitivity detection or with longer bacterial incubation time, are expected to provide more reliable data to examine the robustness of our findings.

Notably, the Pta-AckA pathway and expression of acetyl-CoA synthetase genes (*acs, acsA_1*, *acsA_2*) were suppressed, especially when increased acetate excretion was observed **(Fig. 5C,D,H-J)**. The Pta-AckA pathway is essential for acetate production and metabolic regulation in bacteria (Enjalbert et al., 2017), whereas Acetyl-CoA synthetase catalyzes conversion of acetate to acetyl-CoA (Renilla et al., 2012; Starai & Escalante-Semerena, 2004). Their downregulation suggests a reduction of acetate consumption in 8E, likely contributing to intracellular acetate accumulation and its efflux (Enjalbert et al., 2017; Renilla et al., 2012).

### Thermal sensitivity of DMSP cleavage in *E. ruthgatesiae* 8E

In 8E we found significantly higher DMSP cleavage activity at 25°C than at 31°C. Gardner *et al*. (2022) demonstrated that thermal stress (32°C) significantly increases DMSP concentration in the coral, *Acropora millepora*, as well as the proportion of DMSP being catabolized via the *dddP*-dependent cleavage pathway (Gardner et al., 2022). While this indicates a global metabolic shift toward DMSP cleavage in the coral holobiont at elevated temperatures, our metabolism assays with a pure bacterial culture suggest that the DMSP scavenging capacity of 8E decreases with rising temperatures. Motard-Côté and Kiene (2015) reported that bacterioplankton tend to retain DMSP in cells under high salinity (Motard-Côté & Kiene, 2015), supporting a protective function of DMSP in bacteria (Broy et al., 2015; Cosquer et al., 1999). These findings suggest that microbes under stress may prioritize DMSP accumulation as an anti-stress agent rather than catabolizing it, possibly explaining the reduced DMSP cleavage activity we observed. Nevertheless, as the coral holobiont environment is significantly more complicated than *in vitro* cultures, the ecological role of 8E in response to thermal stress remains to be further investigated.

### DMSP inhibits growth and central carbon metabolism in 8E

At both temperatures, DMSP exposure suppressed the growth of 8E (**Fig. 2A,B**). Whereas no significant decrease in OD_600_ of 8E was observed with 0.1 mM DMSP exposure (Chiou et al., 2023), the reduced OD_600_ here with 3 mM DMSP suggests that DMSP may impose a dose-dependent metabolic burden on 8E. At 25°C, OD_600_ paralleled live cell density at 5 h and 9 h (**Fig. 2A,C**). At 31°C, by contrast, the OD_600_ of the control group was significantly higher than those of the two DMSP-treated groups, whereas live cell density did not differ among groups (**Fig. 2B,D**). Viable cells in 3 mM DMSP-treated group were found smaller than those in the control group at both temperatures (**Fig. S1E,F**). This may explain why the OD_600_ of the control group exceeded those of the DMSP-treated groups at 31°C despite comparable live cell densities.

The growth suppression of 8E by DMSP may be attributed to a direct inhibitory effect of the high DMSP concentration used in this study, or to the accumulation of acetate during DMSP catabolism. Previous research with *E. coli* has shown that acetate accumulates as a byproduct of glucose metabolism and inhibits bacterial growth at high concentrations (De Mey, De Maeseneire, et al., 2007; De Mey, Lequeux, et al., 2007; Pinhal et al., 2019). However, in our experiments, the peak acetate concentration in the medium is associated with the highest OD_600_ value (**Fig. 2B,H**), suggesting that the observed growth suppression may be more associated with cellular toxicity of DMSP.

Genetically, the TCA cycle was suppressed under DMSP exposure (**Fig. 6**). Suppression of the TCA cycle could restrain incorporation of carbon into bacterial biomass, explaining the observed growth suppression and potentially the absence of the ^13^C signal in 8E cells. In addition, *glcB* (malate synthase), *maeA*, and *maeB* (malate dehydrogenase) were slightly upregulated (**Fig. 5E-G)**. These genes are closely linked to metabolism of malate, a key intermediate that allows acetyl-CoA to bypass the decarboxylation process via the glyoxylate shunt (Afzal et al., 2023; Amorim Franco & Blanchard, 2017; Blombach et al., 2007; Huergo et al., 2020; Maloy et al., 1980; Marienhagen & Eggeling, 2008; Renilla et al., 2012). Thus, we hypothesize that high concentrations of DMSP trigger a metabolic shift in 8E, diverting resources from rapid proliferation toward other cellular functions (Ahn et al., 2016; Park et al., 2019).

### DMSP mitigates the genetic response to thermal stress in 8E

Interestingly, while the 31°C control group at 9 h diverged from all other treatments transcriptomically, the DMSP-treated group exhibited a profile resembling those at 25°C (**Fig. 3**). This suggests that DMSP may mitigate transcriptomic alteration of 8E induced by thermal stress. These findings imply that DMSP may bolster 8E’s resistance to thermal stress. In *Ruegeria pomeroyi*, DMSP addition activated DMSP cleavage activity and reduced expression changes in oxidative stress genes (Wang et al., 2022). In symbiotic systems, this potentially allows stabilization of the metabolic state of associated bacteria under stressful conditions. Nevertheless, the distinct transcript profile of 8E in the non-DMSP control group at 31°C, 9 h may also reflect a difference in growth stage (**Fig. 2B**), which was at mid-to-late log phase compared to other groups (at early log phase). This differentiation warrants further study.

In this study, we demonstrate acetate excretion during DMSP cleavage in *Endozoicomonas ruthgatesiae* 8E and provide evidence that it originates from DMSP. Given that acetate is a readily accessible carbon source for most organisms, this DMSP-derived carbon may supply other symbionts in coral holobionts. Transcriptomic data revealed a reconfiguration of metabolic pathways in the presence of DMSP. Specifically, downregulation of the TCA cycle, combined with upregulation of *dddD*, defines a metabolic strategy that prioritizes DMSP processing over biomass production. Furthermore, we observed that elevated temperature diminishes DMSP cleavage activity in 8E. This study offers a new perspective on DMSP metabolism in coral symbiotic bacteria and opens new avenues for future research, in which high-sensitivity isotopic flux analysis and genetic manipulation will be essential to resolve these metabolic dynamics and their ecological implications in coral holobionts.

## Author contributions

**Ying-Chieh Chen**: Conceptualization, Methodology, Software, Formal analysis, Investigation, Data curation, Writing – original draft, Writing – review & editing, Visualization;

**Jui-Hung Yen**: Investigation, Writing – review & editing;

**Ting-Chang Hsu**: Methodology, Investigation, Writing – review & editing;

**Wan-Ting Liao**: Investigation, Writing – review & editing;

**Hsin-Feng Chang**: Investigation, Writing – review & editing;

**Chih-Ying Lu**: Investigation, Writing – review & editing;

**Li-Rong Lin**: Investigation, Writing – review & editing;

**Po-Shun Chuang**: Conceptualization, Methodology, Supervision, Project administration, Funding acquisition, Writing – review & editing;

**Sen-Lin Tang**: Conceptualization, Resources, Supervision, Project administration, Funding acquisition, Writing – review & editing.

## Acknowledgements

We thank Dr. Yu-Fang Tseng from the Comprehensive Flow Cytometry Laboratory, Dr. Yu-Ching Wu from the Small Molecule Metabolomics Core Laboratory, and Dr. Sung-Yun Hsiao and Dr. Der-Chuen Lee from the NanoSIMS Laboratory of Academia Sinica for their technical support. We thank Dr. Steven D. Aird for editing and commenting on this manuscript. The authors used Gemini 3 and Claude Opus 5.5 for language editing during the writing of this manuscript. All final interpretations and conclusions were formulated by the authors, who take full responsibility for the integrity and accuracy of the work.

