## Supplementart information for "Integrated transcriptomic and metabolite analyses link DMSP cleavage to acetate overflow in the coral symbiont *Endozoicomonas ruthgatesiae*"

**Table S1. ANOVA between different temperatures and timepoints (corresponding to Figure 2).**

| | Temperature (°C) | Timepoint (h) | *p* value |
|---|---|---|---|
| $OD_{600}$ | 25 | 0 | 0.5787 |
| | | 5 | 0.0300 |
| | | 9 | 0.0079 |
| | 31 | 0 | 0.2160 |
| | | 5 | 0.0006 |
| | | 9 | 2.44E-05 |
| DMS | 25 | 0 | 0.3377 |
| | | 5 | 9.12E-09 |
| | | 9 | 5.18E-08 |
| | 31 | 0 | 0.4219 |
| | | 5 | 6.15E-06 |
| | | 9 | 0.0074 |
| Acetate | 25 | 0 | 0.3569 |
| | | 5 | 0.6431 |
| | | 9 | 0.0207 |
| | 31 | 0 | 0.3855 |
| | | 5 | 0.0050 |
| | | 9 | 0.0028 |
| Live cell density (cells $mL^{-1}$) | 25 | 5 | 0.0316 |
| | | 9 | 0.0081 |
| | 31 | 5 | 0.5291 |
| | | 9 | 0.1146 |
| Acetate production per cell (µmol $cell^{-1}$) | 25 | 5 | 0.0515 |
| | | 9 | 0.0024 |
| | 31 | 5 | 0.0035 |
| | | 9 | 0.0089 |

**Table S2. DEGs that overlapped different temperatures and timepoints.**

| Regulation | Region (temperature[°C]_timepoint[h]) | Gene display |
|---|---|---|
| Up-regulated | 25_5 & 25_9 | hypothetical protein |
| | 25_5 & 25_9 & 31_5 & 31_9 | *dddD* |
| | 25_5 & 25_9 & 31_5 & 31_9 | hypothetical protein |
| | 25_5 & 25_9 & 31_5 & 31_9 | hypothetical protein |
| | 25_5 & 25_9 & 31_9 | *grxA* |
| | 25_5 & 25_9 & 31_9 | hypothetical protein |
| | 25_5 & 25_9 & 31_9 | hypothetical protein |
| | 25_5 & 25_9 & 31_9 | hypothetical protein |
| | 25_5 & 31_9 | *gpx1* |
| | 25_5 & 31_9 | *tadA_3* |
| | 25_5 & 31_9 | *yxeM* |
| | 25_5 & 31_9 | hypothetical protein |
| | 25_5 & 31_9 | hypothetical protein |
| | 25_9 & 31_5 & 31_9 | hypothetical protein |
| | 25_9 & 31_9 | *fdxA* |
| | 25_9 & 31_9 | *yciO* |
| | 25_9 & 31_9 | *yfgD* |
| | 25_9 & 31_9 | hypothetical protein |
| | 25_9 & 31_9 | hypothetical protein |
| | 25_9 & 31_9 | hypothetical protein |
| | 25_9 & 31_9 | hypothetical protein |
| | 25_9 & 31_9 | hypothetical protein |
| | 25_9 & 31_9 | hypothetical protein |
| | 25_9 & 31_9 | hypothetical protein |
| | 25_9 & 31_9 | hypothetical protein |
| | 25_9 & 31_9 | hypothetical protein |
| | 25_9 & 31_9 | hypothetical protein |
| Down-regulated | 25_5 & 25_9 | hypothetical protein |
| | 25_5 & 25_9 | hypothetical protein |
| | 25_5 & 25_9 | hypothetical protein |
| | 25_5 & 25_9 | hypothetical protein |
| | 25_5 & 25_9 | hypothetical protein |
| | 25_5 & 25_9 | hypothetical protein |
| | 25_5 & 25_9 & 31_5 | *pilE1* |
| | 25_5 & 25_9 & 31_5 | *ousW* |
| | 25_5 & 25_9 & 31_5 | *proV* |
| | 25_5 & 25_9 & 31_5 | *xdhA* |
| | 25_5 & 25_9 & 31_5 | hypothetical protein |
| | 25_5 & 25_9 & 31_5 | hypothetical protein |
| | 25_5 & 25_9 & 31_9 | *gbpA* |
| | 25_5 & 25_9 & 31_9 | *oprD* |
| | 25_5 & 25_9 & 31_9 | hypothetical protein |
| | 25_5 & 25_9 & 31_9 | hypothetical protein |
| | 25_5 & 31_5 | hypothetical protein |
| | 25_5 & 31_5 & 31_9 | hypothetical protein |
| | 25_5 & 31_9 | hypothetical protein |
| | 25_5 & 31_9 | hypothetical protein |
| | 25_5 & 31_9 | hypothetical protein |
| | 25_5 & 31_9 | hypothetical protein |
| | 25_5 & 31_9 | hypothetical protein |
| | 25_9 & 31_5 | *fklB_2* |
| | 25_9 & 31_5 | hypothetical protein |
| | 25_9 & 31_5 | hypothetical protein |
| | 25_9 & 31_9 | *hxuB* |
| | 25_9 & 31_9 | *trpD* |
| | 25_9 & 31_9 | *dmdC_2* |
| | 25_9 & 31_9 | hypothetical protein |
| | 31_5 & 31_9 | hypothetical protein |
| | 31_5 & 31_9 | hypothetical protein |

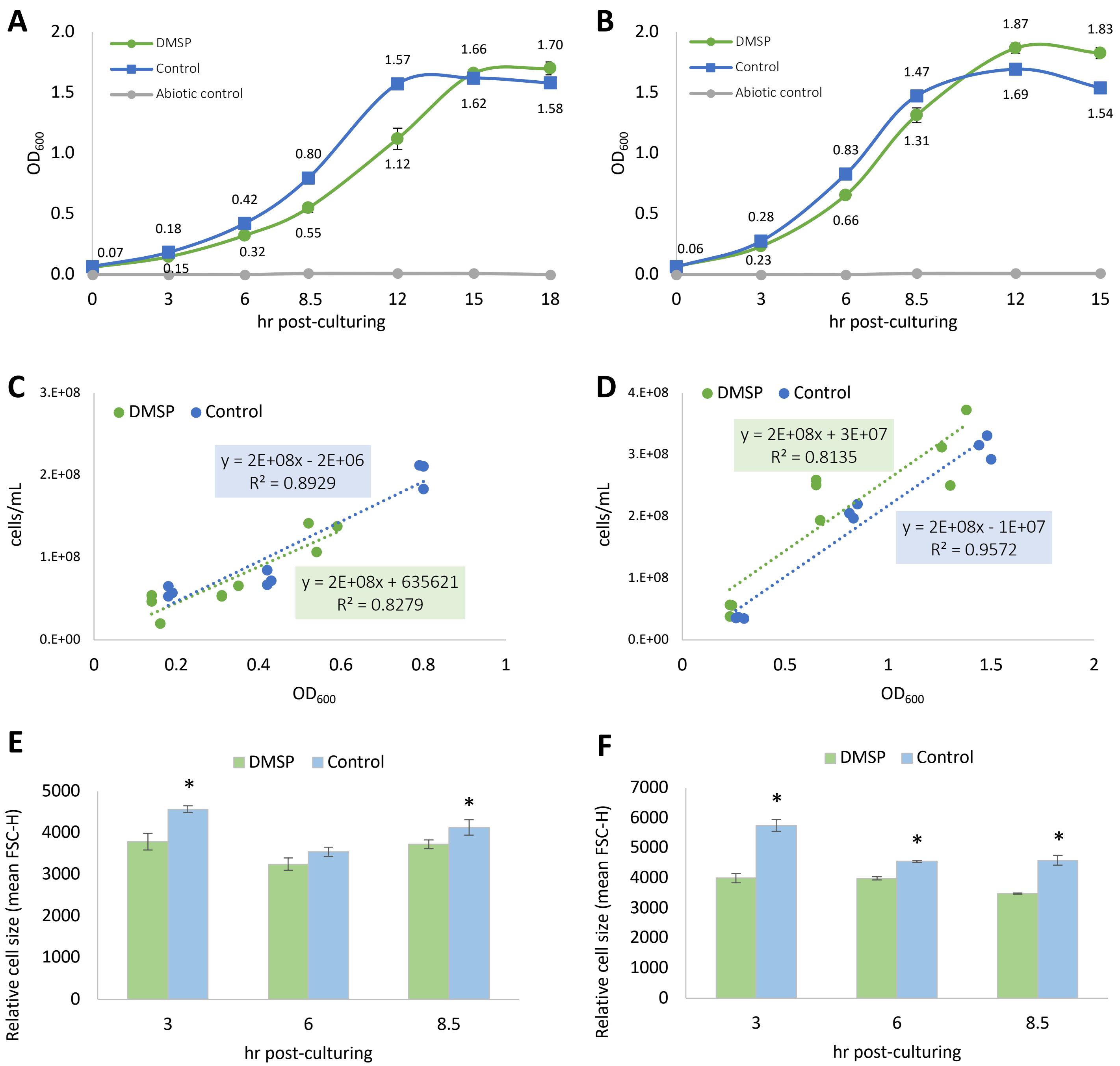


**Figure S1. Growth curves and linear regressions for optical density [600 nm ($OD_{600}$)] and live cell density of *Endozoicomonas ruthgatesiae* 8E under different conditions.** An independent set of 300-mL sterile, gas-tight vials from the DMSP metabolism assay and RNA sequencing was cultured to generate growth curves and to establish $OD_{600}$-to-cell density conversions. Vials were incubated at (A) 25°C and (B) 31°C to evaluate temperature-dependent growth. Culture aliquots were sampled using the same method as employed in the DMSP metabolism assay at 0, 3, 6, 8.5, 12, 15 h. Linear regression between $OD_{600}$ and live cell density (cells $mL^{-1}$) at (C) 25°C and (D) 31°C was measured at 3, 6, 8.5 h, and regression lines of different treatment-time group were established. Mean forward scatter height (FSC-H) of live cells at (E) 25°C and (F) 31°C, measured by flow cytometry as a relative proxy for cell size. The DMSP and control groups in (A,B) each had three biological replicates (n=3), and an abiotic (uninoculated, n=1) control was included to confirm the absence of medium contamination; in (C–F), the DMSP and control groups each comprised three biological replicates, each measured in two technical replicates.

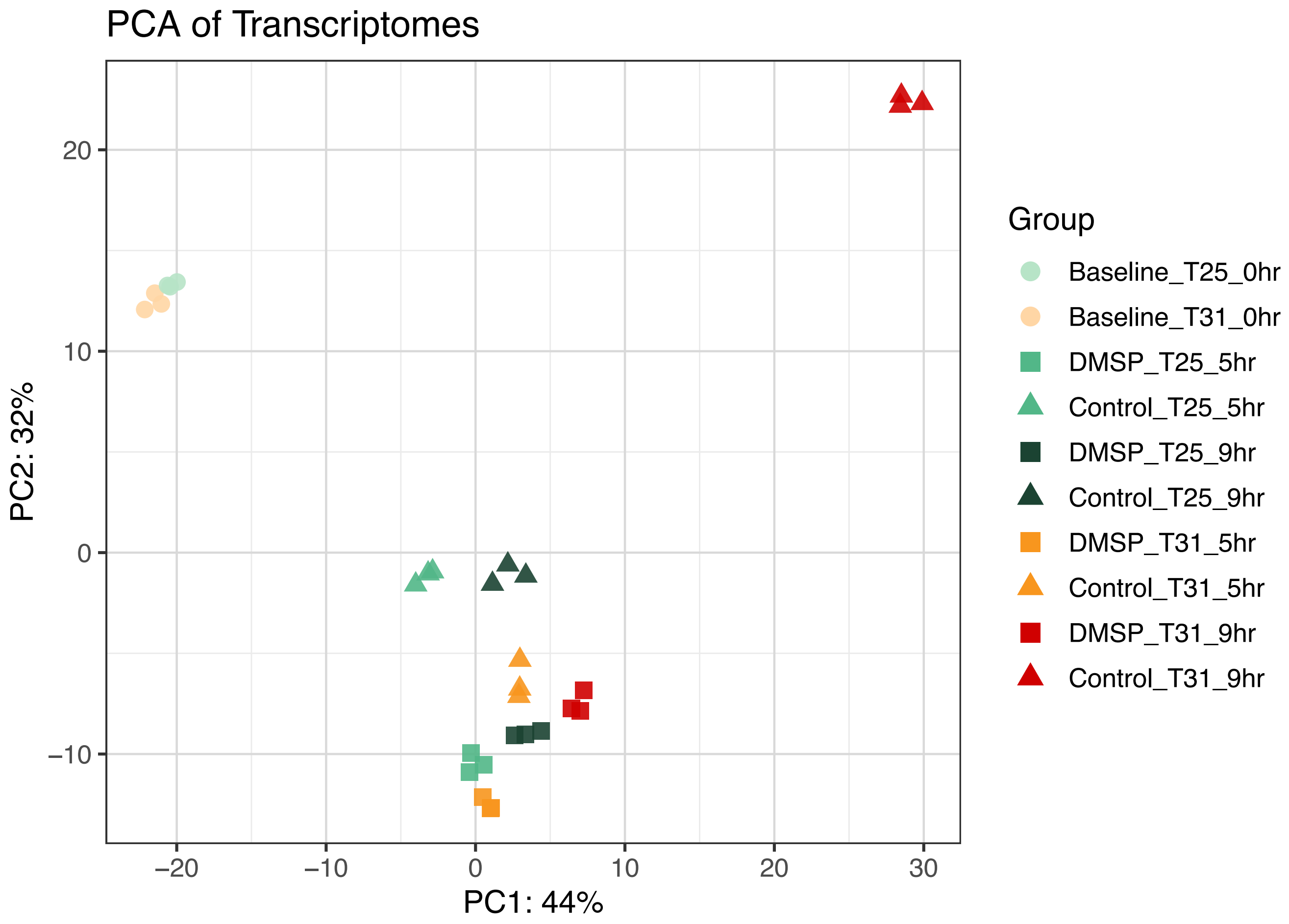


**Figure S2. Principal components analysis (PCA) of 8E transcript profiles under different treatments and timepoints, including baseline (0 h).** This analysis used the same method employed in **Fig. 3**, with the baseline included, representing the 0 h timepoint of the culture initially grown in MMBv4 prior to centrifugation, washing, and transfer into minimal medium. Baseline transcriptomic profiles did not differ significantly between the 25°C and 31°C experiments (PERMANOVA, $R^2 = 0.52$, $p = 0.10$. Group dispersions also did not differ, PERMDISP $p = 0.70$), although statistical power was limited by the small sample size (n = 3 per temperature).
